# Microbially induced calcite precipitation using *Bacillus velezensis* with guar gum

**DOI:** 10.1101/634212

**Authors:** Rashmi Dikshit, Animesh Jain, Arjun Dey, Sujit Kamilya, Abhishake Mondal, Aloke Kumar

**Affiliations:** Dept. of Mechanical Engineering, Indian Institute of Science, Bangalore-560012, India; Thermal Systems Group, U. R. Rao Satellite Centre (Formerly known ISRO Satellite Centre), Indian Space Research Organisation, Bangalore-560017, India; Solid State and Structural Chemistry Unit, Indian Institute of Science Bangalore-560012, India

**Keywords:** microbially induced calcite precipitation, *Bacillus velezensis*, guar gum, biocementation, ureolysis

## Abstract

The present study was performed to explore the efficiency of microbially induced calcite precipitation (MICP) via locally isolated bacterial strains. Strains were isolated from soil and were screened for urease activity as well as microbial precipitation. Among all screened isolates, a carbonate precipitating soil bacterium was subjected for 16S rRNA gene sequencing. This strain was identified as *Bacillus velezensis*. The MICP characteristics of this strain were explored under three different media compositions and significant amount of precipitation in all cases was observed. Highest amount of precipitation was seen with guar as a biopolymer additive medium under experimented conditions. Activity of isolated strain with reference to pH profile, and ammonia concentration and total reducing sugar was further explored under media supplemented with four concentrations of guar (0.25%, 0.5%, 0.75% and 1% w/v). Microstructural analysis of microbial precipitation was performed with the help of scanning electron microscopy (SEM) and X-ray powder diffraction (XRD) analysis, which confirmed the presence of calcium carbonate in different phases. The strain was subjected to bio-cementation and locally available sand was successfully consolidated. XRD results confirmed the presence of calcium carbonate on consolidated samples.

## 1. Introduction

Mineral precipitation mediated via microbial metabolic activity known as bio-mineralization is a pervasive phenomenon on our planet. Reactions such as sulfate reduction [1], methane oxidation [2], photosynthesis [3], and urea hydrolysis [4] help in either increasing the environmental pH or metabolization of dissolved inorganic carbon [5]. Microbial induced calcium carbonate precipitation (MICP), one of the mechanisms of bio-mineralization, is a process which creates a favourable microenvironment for calcium carbonate precipitation by microbes [6] MICP has potential applications in numerous domains such as prevention of soil erosion [7, 8], enhancing the durability of concrete [9], restorations of cultural and historical assets [10–12] and production of bio-cement [13, 14].

Urea hydrolysis is an important pathway for MICP due to its exclusively controlled mechanism to get higher concentration of precipitated carbonates [15]. Diverse bacterial strains such as *Sporosarcina pasteurii* (formerly *Bacillus pasteurii*)[16, 17], *Bacillus megaterium* [14]*, Helicobacter pylori* [16]*, Pseudomonas aeruginosa* [16], etc are known to hydrolyse urea by producing the urease enzyme (urea amidohydrolase; EC 3.5.1.5). *Sporosarcina pasteurii*, an alkaliphilic, non-pathogenic and endospore producing bacteria has been extensively explored for MICP [18–21]. Achal et al. [19] developed a mutant strain of *S. pasteurii* to enhance urease activity which has shown significantly more urease activity and CaCO_3_ precipitation compared to the wild strain of *Sporosarcina pasteurii MTCC 1761*. Dhami et al. [22] reported improved concrete strength and durability of construction material when incubated with *Bacillus megaterium*. Strains belonging to *Bacillus* genus are suitable for MICP applications as they can tolerate adverse environmental stresses such as alkaline pH, higher mechanical forces and dehydrating conditions by forming endospores [5, 23]. Given the relevance of MICP to various applications, discovery of new naturally occuring MICP-capable strains is likely to be beneficial. In this study, an unexplored soil isolated *Bacillus* strain namely *Bacillus velezenesis* was explored for MICP.

Moreover, in nature, biominerals are mainly embedded in organic matrices, made up of macromolecules. These macromolecules can be proteins or polysaccharides, known to act as templates for growth and precipitation of bio-minerals[24, 25]. Organic macromolecules along with inorganic minerals have proven to be nucleating sites for mineral precipitation[26–28]. Also, catanionic polypeptides influence CaCO_3_ precipitation as it contains negatively charged carboxylic group, which shows strong affinity towards Ca^2+^ in the presence of CO_3_^2−^ to promote CaCO_3_ nucleus formation[28]. Here we explored the role of two biopolymers on the MICP process: a) the guar and b) L-phenylalanine, which is a cationic polypeptide. Guar (*Cyamopsistetragonolobus*, family Leguminosae) is a green, non-toxic, easily available, cost-effective and easily biodegradable polysaccharide-based natural polymer[29]

The aim of present study was to isolate and characterize indigenous strains for calcium carbonate precipitation under three different media compositions and its application to soil consolidation. This was accomplished by (i) screening of soil isolated bacterial strains for urease activity and precipitation of CaCO_3_ via ureolytic pathway, (ii) exploration of precipitation under various nutritional conditions, (iii) investigation of growth kinetics of urease positive strain with different concentration of guar gum supplementations and (iv) consolidation of locally available sand in order to check bio-consolidation efficacy of indigenous isolated strain(s).

## 2. Materials and Methods

### 2.1 Bacterial strains and its characterization

Soil isolated bacterial strains used for the present study were gratefully obtained from Dr. Swetha S. and maintained on glycerol stock at −80°C. Strains were sub cultured on nutrient agar media at 30°C for 24 hours. Seed culture was prepared by inoculating one bacterial colony with 50 ml nutrient broth media in shaking incubator (BioBee, India) at 120 rpm and 30°C to get optical density of 0.5 at 600 nm. The strains were named as SI1 (Soil isolate 1), SI2 (Soil isolate 2), SI3 (Soil isolate 3) and SI4 (Soil isolate 4). Biochemical characterization of isolated strains was done by performing biochemical assay such as oxidation-fermentation test, catalase test, Voges-Proskauer test, sugar fermentation, etc with the help of *Bacillus* identification kit (Hi-media, India).

### 2.2 Screening for Urea hydrolysis

Isolated strains were screened for urea hydrolysis by streaking a single bacterial colony on urea-agar plate and kept at 30°C for 24 hours. To observe change in medium pH, phenol red was added as an indicator along with media. Urea-agar plate was prepared by adding 0.1 g glucose, 0.1 g peptone, 0.5 g NaCl, 0.2 g mono-potassium phosphate, 2 g urea, 0.0012 g phenol red dye and 2 g agar to 100 ml distilled water. All chemicals were procured from Hi-Media, India.

### 2.3 Molecular identification of isolate

Molecular identification of urease positive strain was done by 16S rRNA gene sequencing. Colonies from single streak on the agar plate were scraped and suspended in PBS buffer and centrifuged. The pellet obtained was dispersed in 600 µl of cell lysis buffer (Guanidium isothiocyanate, SDS, Tris-EDTA) and mixed by inverting the vial for 5 minutes and incubated for 10 minutes with gentle mixing till the suspension looked almost transparent. The solution was layered on top with 600 µl of isopropanol. The two layers were mixed gently till white strands of DNA were seen and the solution became homogenous. The spooled DNA was spun to precipitate DNA at 10,000 rpm for 10 minutes. The air-dried pellet was suspended into 50 µl of 1X Tris-EDTA buffer and incubated for 5 min at 55–60°C to increase the solubility of genomic DNA. 5 µl of freshly extracted DNA along with 3µl of gel loading dye was loaded onto 1% agarose gel and subjected to electrophoresis. Amplification of 16s rRNA gene was performed using the following primers.

Forward primer: 5’-AGAGTTTGATCCTGGCTCAG-3’

Reverse primer: 5’-ACGGCTACCTTGTTACGACTT-3’

### 2.4 In vitro CaCO_3_ Precipitation

Urease positive strain was evaluated for microbial induced precipitation under flask conditions in the following media compositions:

1. Synthetic media-urea-calcium chloride (hereafter, SMUC): Prepared by adding 0.1 g glucose, 0.1 g peptone, 0.5 g NaCl, 0.2 g mono-potassium phosphate and 2 g urea in 100 ml of distilled water with 25 mM CaCl_2_.
2. Synthetic media-urea-calcium chloride-guar gum (hereafter, SMUCG): Prepared by replacing glucose in SMUC medium with 1% (w/v) guar gum (Urban Platter, India).
3. Synthetic media-urea-calcium chloride-L-phenylalanine (hereafter, SMUCP): Prepared by adding 50 mg/l L-phenylalanine (Hi-Media, India) in SMUC medium.

The strain was inoculated in all mentioned media and incubated at 30°C for 7 days under static condition. After the incubation period, samples were centrifuged (Sorvall^TM^ Legend^TM^ X1 Centrifuge, Thermo Fisher Scientific, Germany) at 4°C and 5000 rpm for 10 minutes. The supernatant was discarded and precipitates were dried at 37°C for 12 hours in hot air oven (BioBee, India). Precipitates were weighed using analytical weighing balance (ATX224 Unibloc®, Shimadzu, Japan). These precipitates were further observed using scanning electron microscope imaging and X-ray diffraction analysis.

### 2.5 Bacterial activity with guar gum

Bacterial activity of isolated urease positive strain was investigated in SMUCG medium supplemented with varying concentration of guar gum (0.25%, 0.50%, 0.75% and 1% w/v) and was compared with SMUC medium. The strain was inoculated and incubated at 30°C, pH (CyberScan pH meter, Eutech Instruments), ammonium concentration and amount of reducing sugar were measured at different time intervals. Nessler’s reagent assay was used to quantify ammonium concentration, by adding 100 µl of Nessler’s reagent (Hi-Media, India) in 2 ml of bacterial sample. Absorbance was measured at 425 nm (using UV/Vis spectrophotometer, Shimadzu, Japan) after 10 minutes of incubation [30]. Total reducing sugar was estimated using DNS protocol [31] by measuring optical density at 540 nm.

### 2.6 Preparation of sample for bio-consolidation

A 50 ml Disposable polypropylene syringes (DispoVan^TM^) with an internal diameter of 30 mm were used as a bioreactor for consolidation of loose mass (Fig 1). The outlet of syringe was sealed using parafilm and inner surface was layered with OHP transparency film to ensure easy removal of sample. The column was fitted with scouring pad at bottom and packed with locally available manufactured sand (M-sand) (particle size characteristics: d_10_(10% of sand particles have this diameter or lower) = 125 µm; d_50_ = 212 µm; d_90_ = 250 µm) and covered again with a layer of scouring pad. To avoid influence of any external contamination, sand was autoclaved at 121°C and 15 psi for 30 minutes. The top portion of bioreactor was kept empty for aeration and was covered with cotton plug to avoid contamination.

**Fig 1:**
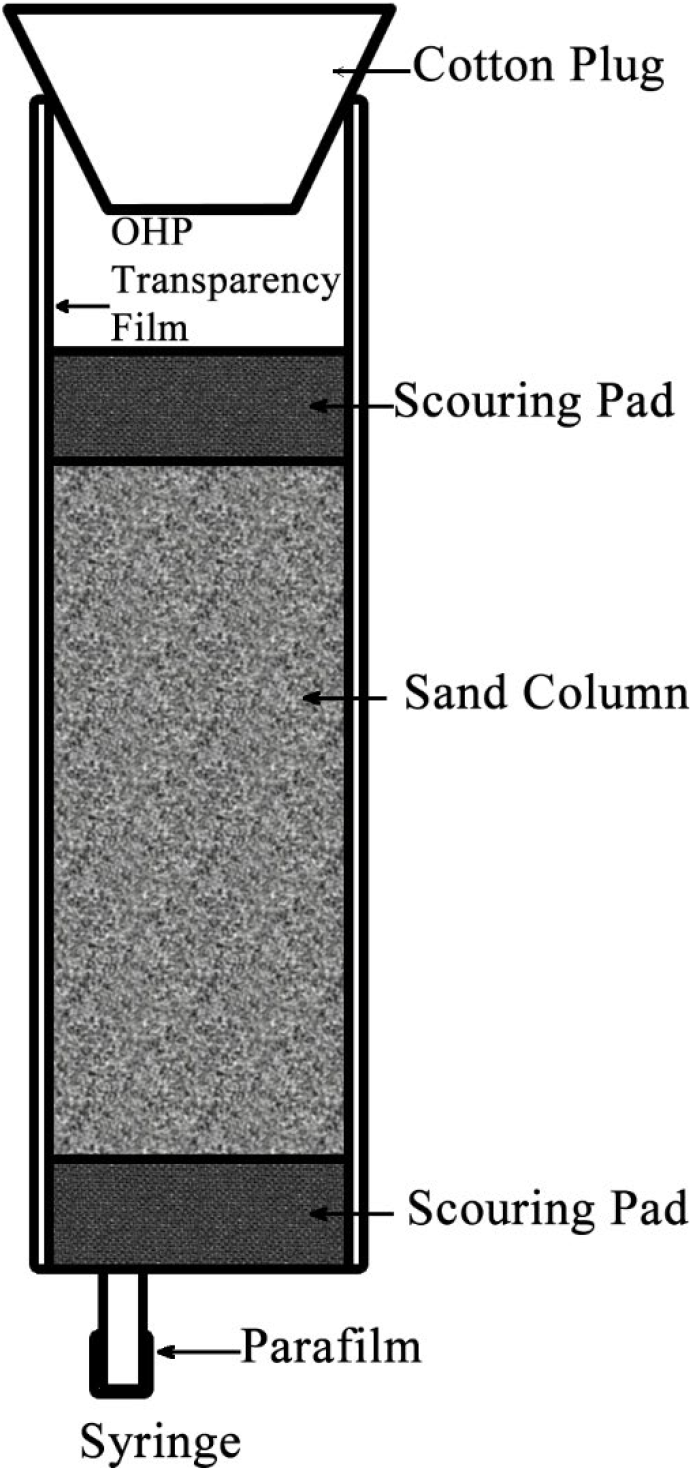
Schematic of Experimental Setup

Inoculum was prepared by culturing *Bacillus velezensis* in SMUC media at 30°C and 130 rpm for 24 hours to obtain an optical density of 2 at 600 nm (OD_600_). A volume of 5 ml inoculum was added to sand column along with 5 ml of SMUC media and column was maintained at 30°C. After 72 hours, extra culture at top of column was pipetted out and 5 ml SMUC media along with 5 ml of bacterial culture was injected from bottom of the syringe to ensure flow in every portion of column. This process was repeated after 144 hours. Sample was removed after 240 hours and kept for drying at 45°C for 24 hours. The experiments were performed in triplicates. The result was compared with two control samples: (a) sand with tap water and (b) sand with SMUC media (without bacteria) under same treatment.

### 2.7 Scanning electron microscopy (SEM)

Bacterial precipitates in different media compositions (SMUC, SMUCG and SMUCP) and bio-consolidated samples were observed using Scanning Electron Microscopy (Carl Zeiss AG-ULTRA 55, Germany). The dried precipitates were fixed with 2.5% glutaraldehyde in PBS buffer (pH 7.4) for 30 minutes, followed by centrifugation at 5°C and 5000 rpm for 5 minutes. The supernatant was removed and sample was washed with PBS buffer to remove fixative. The treated samples were dehydrated with serial dilution using 10, 30, 50, 70, 90 and 100% v/v ethanol. The samples were mounted on carbon tape fixed on aluminium stub and kept in desiccator for 24 hours. Before SEM imaging, the dried samples were coated with gold for 120sec to avoid charging.

### 2.8 X-ray diffraction (XRD) analysis

The precipitated samples and bio-consolidated sand were collected, air dried and grinded to fine powder. The fine powder was analysed using a commercial X-ray diffractometer (powder XRD, Rigaku SmartLab®, Rigaku Corporation) and thoroughly indexed as per Inorganic Crystal Structure Database (ICSD) library using PANalytical X’PertHighScore Plus pattern analysis software.

## 3. Results

### 3.1 Screening and identification of strains

Morphological and biochemical characterisation of all soil isolated strains was performed with the help of *Bacillus* kit. All strains were found to be gram-positive, aerobic, spore-forming and catalase-positive (Table S1). Further, qualitative analysis based on pH indicator (phenol red) for urea degradation of all strains was performed on SMU agar plate (Fig 2). Colour change was observed only in case of SI1 (Fig 2a) due to enhancement in medium pH while no change was observed in SI2 (Fig 2b), SI3 (Fig 2c) and SI4 (Fig 2d) as compared to control (without organism) (Fig 2e). SI1 was subjected to 16S rRNA gene sequencing (Fig 3) and based on DNA−DNA relatedness values has shown approximately 99% similarity with *Bacillus amyloliquefaciens* and was identified as *Bacillus velezensis* (Table S2).

**Fig 2:**
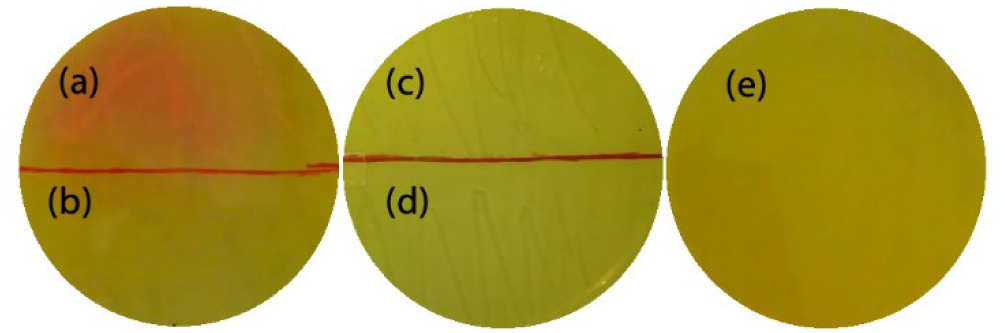
Plate showing urea hydrolysis after 24hrs of Soil Isolated strains (a) SI1, (b) SI2, (c) SI3, (d) SI4, (e) Control (without organism). SI1 shows color change due to increase in medium pH by bacterial activity

**Fig 3:**
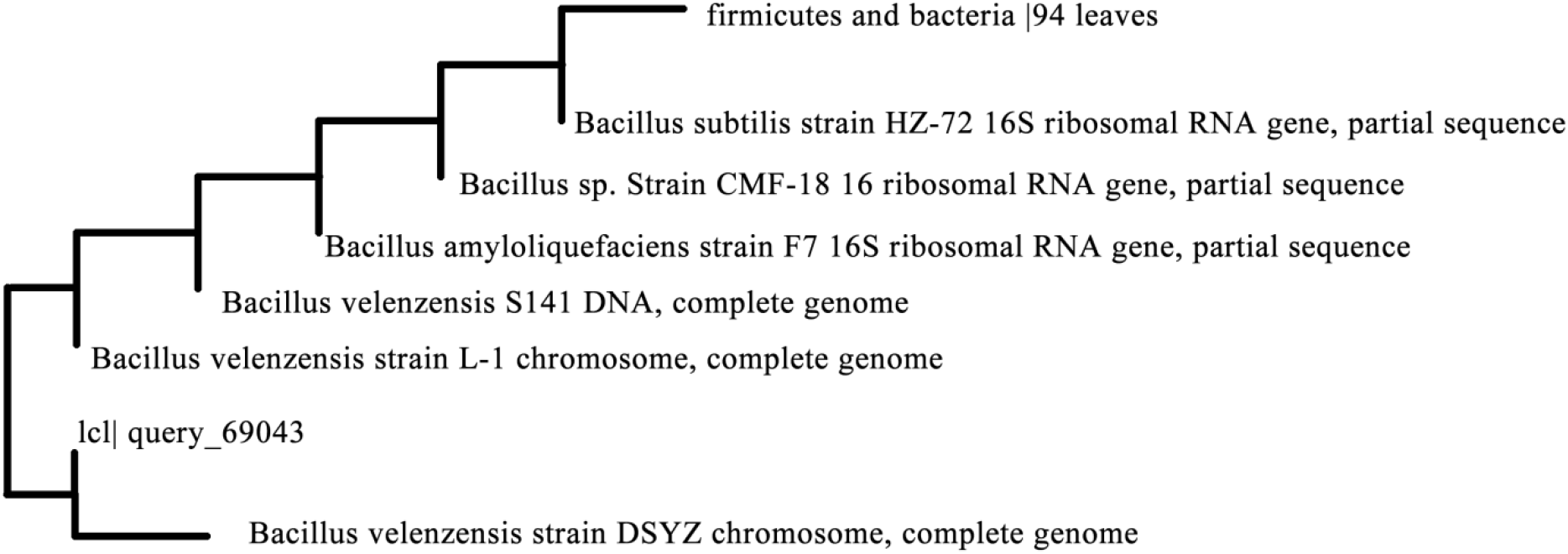
Phylogenetic tree of 16s rRNA gene sequencing for urease positive strain

### 3.2 Precipitation under flask condition and its microstructural analysis

*Bacillus velezensis* induced precipitates were obtained with three different media supplementations SMUC, SMUCP and SMUCG after 7 days of incubation under flask conditions. Amount of precipitation (g/l) was plotted for different media supplementations (Fig 4a). It was observed that an increase of 101% in SMUCG and 58% in SMUCP media took place as compared to SMUC medium. This indicates the two additives support bacterial activity resulting in enhanced amount of precipitation.

**Fig 4:**
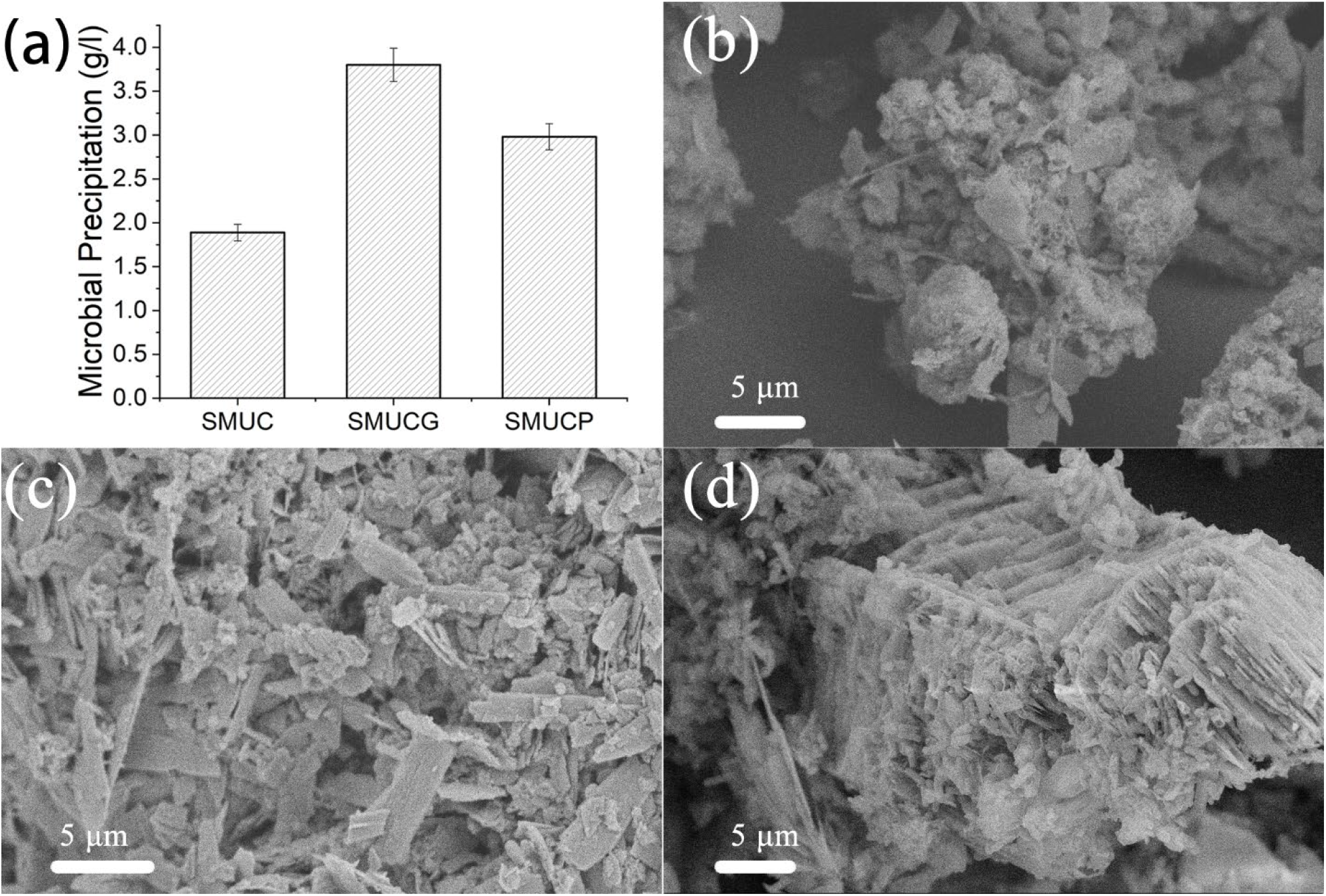
(a) Quantification of microbial precipitation. SEM images of *Bacillus velezensis* mediated precipitated crystals with (b) SMUC medium, (c) SMUCG medium and (d) SMUCP medium. Incubation period was 7 days in all cases.

SEM images of these precipitates with different media supplementations show distinctly different crystal morphologies (Fig 4b,4c,4d). In SMUC medium, sparingly populated mixture of spherical and rectangular shaped crystals embedded in bacterial bioproduct can be observed in Fig 4b. Multi-shaped densely populated sharp pointed flaked precipitates dominated by rectangular flakes (with traces of organic matrix in between) can be observed in Fig 4c in case of SMUCG supplemented media. Whereas, in case of SMUCP medium, the precipitate can be seen to be concentrated in multi-layered stepped plateau like structure with bacteria and its secreted matrix sandwiched in between the layers (Fig 4d).

XRD analysis of dried and crushed precipitated crystals (Fig 5) clearly suggests that different amount of calcite, vaterite and aragonite are co-existing in the samples. The hexagonal vaterite (ICSD File no. 98-011-5332) and orthorhombic vaterite (ICSD File no. 98-010-9797) were found in all three media. SMUC has not shown any calcite content. The identified hexagonal shaped calcite (ICSD File no. 98-002-1912) peaks in SMUCG correspond to 2θ values of 29.15°, 31.44°, 40.72°, 46.57°, 49.52° correlated with lattice indices (hkl) of (1 0 4), (0 0 6), (1 1 3), (0 1 8) and (1 1 6) respectively. While in case of SMUCP, hexagonal shaped calcite (ICSD File no. 98-003-4957) correspond to 2θ values of 23.07°, 29.42°, 31.52°, 36.02°, 39.47°, 43.20°, 47.52°, 48.53°, 57.47° correlated with lattice indices (h k l) of (0 1 2), (1 0 4), (0 0 6), (1 1 0), (1 1 3), (2 0 2), (0 1 8), (1 1 6) and (1 2 2) respectively.

**Fig 5:**
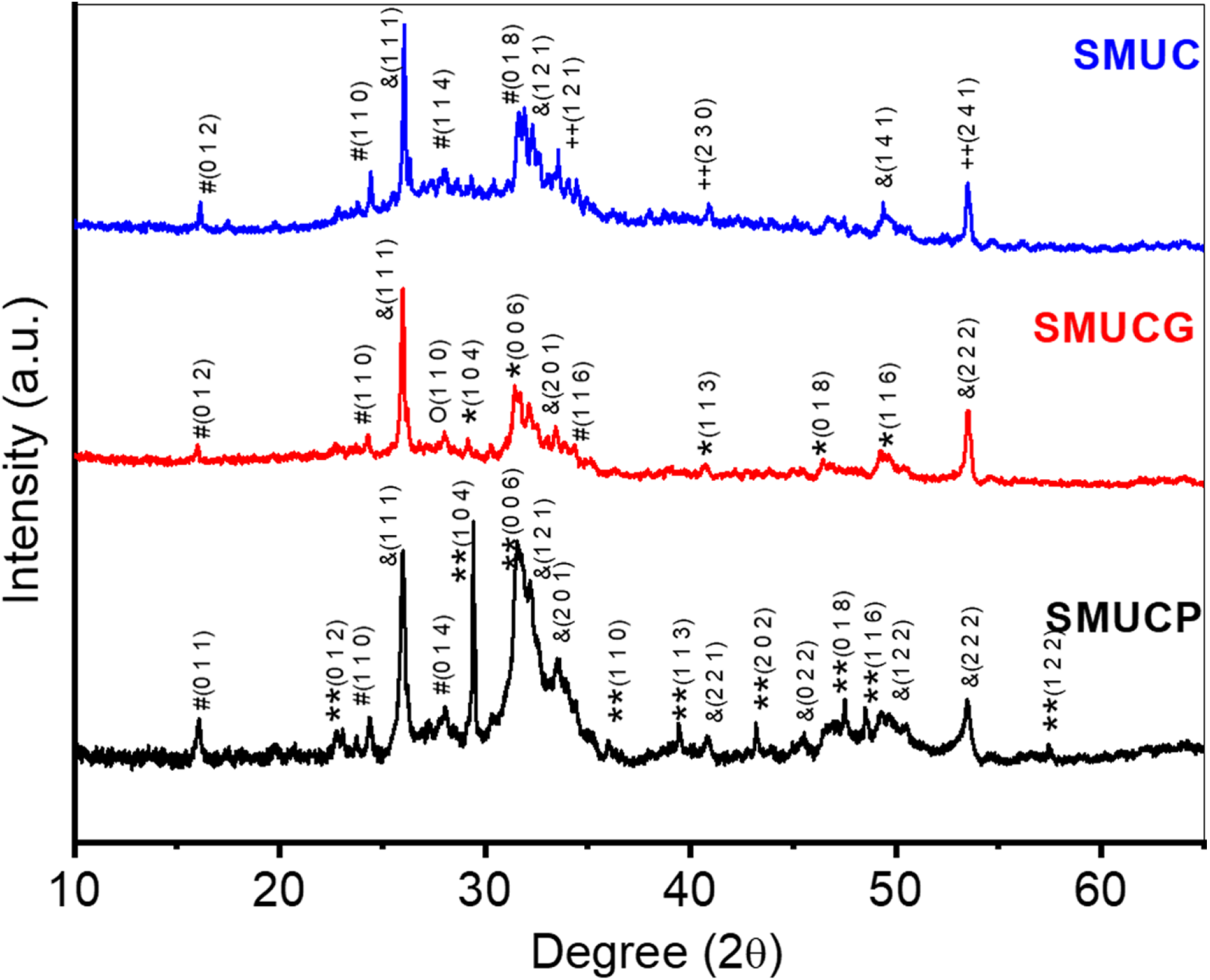
XRD pattern of *Bacillus velezensis* induced precipitates under different media compositions (a) SMUC (b) SMUCG (c) SMUCP. Incubation period was 7 days in all cases. Symbols represent: # (Vaterite ICPD no. 980115332), & (Vaterite ICPD no. 980109797), ++ (Vaterite ICPD no. 980012463), O (Aragonite ICPD no. 980114648), * (Calcite ICPD no. 980021912), ** (Calcite ICPD no. 980034957)

XRD and SEM analysis clearly reveal that different phases of calcium carbonate are precipitated as a result of difference in bacterial activity in all three media. As significant improvement in precipitation was observed in guar gum as a bio-polymer additive medium (SMUCG), its effect was further explored at different guar concentrations.

### 3.4 Influence of polymer on microbial activity and precipitation

*Bacillus velezensis* activity was studied with different concentrations of guar gum (0%, 0.25%, 0.50%, 0.75% and 1% w/v) in SMUCG media. Fig 6a shows the plot of ammonium concentration with time. An increase in ammonium concentration was observed until 96 hours in all cases. Further, it decreased until 144 hours and remained approximately constant thereafter. Maximum ammonium concentration was observed with 1% guar gum additive (17.5 µg/ml) followed by 14.9 µg/ml, 13.5 µg/ml, 11.6 µg/ml, 10.3 µg/ml for 0.75%, 0.50%, 0.25% and 0% guar gum additive respectively. It suggests that an increase in guar gum concentration enhances the rate of urea hydrolysis. This is confirmed by the plot of change in pH of growth medium with time shown in Fig 6a (inset). The experiment was initiated at an acidic pH of 5.5 due to the fact that the media was based on urea and CaCl_2_ composition and any adjustment in pH towards alkalinity may result in quick abiotic precipitation which is undesirable [6]. An increase in pH is observed until 48 hours in all cases while with higher guar gum concentrations, pH further increased until 96 hours. In the case of 0% and 0.25% guar gum supplemented media, pH reaches to maximum value of 8, whereas, a maximum value of 9 was observed in media supplemented with 0.50%, 0.75% and 1% guar gum.

**Fig 6:**
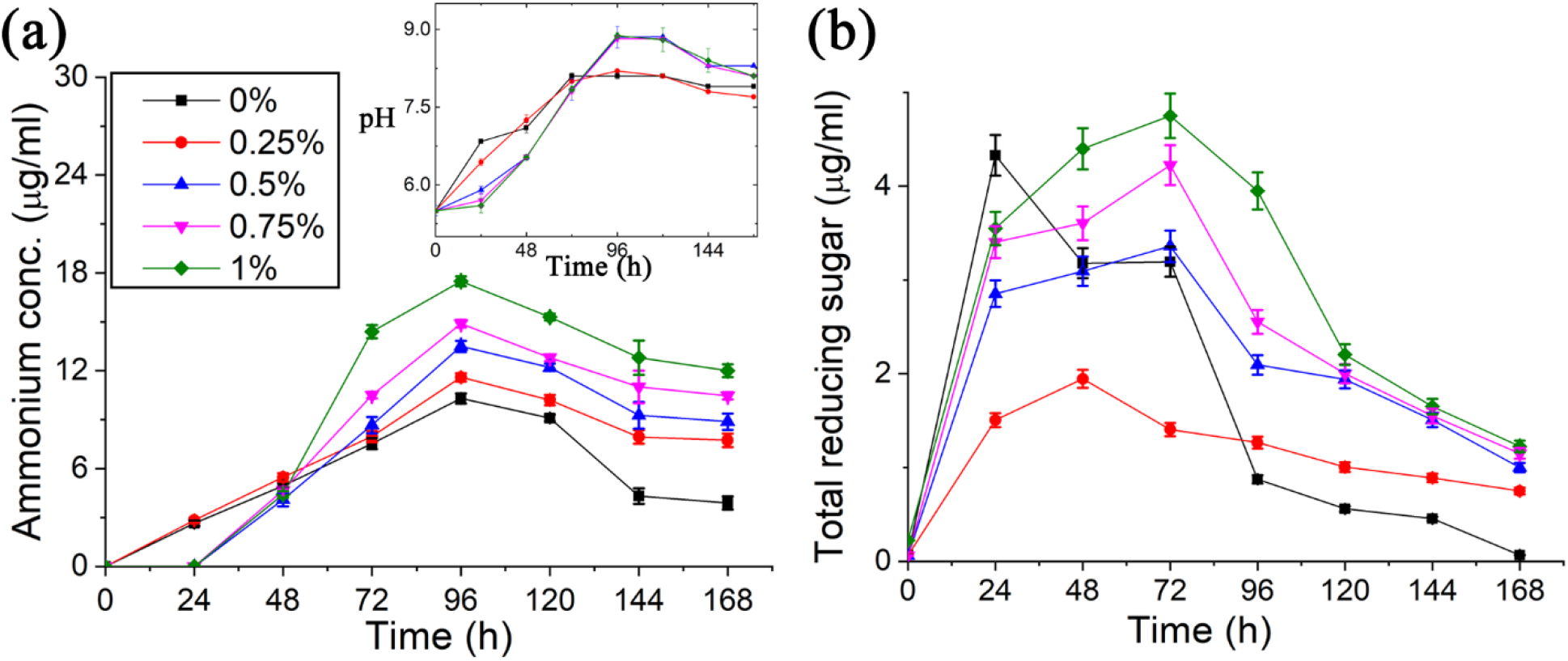
Exploration of microbial physiology under flask condition with guar gum additive. (a) temporal evolution of ammonium concentration in SMUCG media and temporal evolution of pH in SMUCG media (inset) (b) change in amount of total reducing sugar in SMUCG media with time compared with SMUC medium (0% curve). Legend shows guar gum percentage (w/v) in SMUCG media (Error bars represents the standard deviation of the data of three independent experiments).

The effect of guar gum degradation by *Bacillus velezensis* was studied using DNS protocol. In Fig 6b, total reducing sugar is plotted as a function of time. Total reducing sugar content in SMUC media increases till 24 hours followed by its decrease. Interestingly, trends for SMUCG media differed from the control sample (SMUC media). For SMUCG media, amount of total reducing sugar in media increases till 48 hours in 0.25% guar gum while it increases till 72 hours in all other guar gum concentrations and decreases thereafter.

### 3.6 Bio-consolidation of sand

Potential of *Bacillus velezensis* for MICP mediated consolidation was checked in M-sand incubated in SMUC medium. Fig 7a shows consolidated sample of M-sand after 10 days of incubation in bioreactor inoculated with bacterial strain. Control sample with SMUC media (without bacteria) showed no sign of particle aggregation (Fig 7b). Consolidation via bacterial activity was confirmed with SEM imaging and XRD analysis. A bacterial induced matrix bridging sand particles can be observed in SEM image of consolidated sample (Fig 7c) as compared to control sample (Fig 7d). Different sized precipitates can be seen embedded in the matrix of bio-consolidated sample. XRD analysis of bio-consolidated sample when compared to control (Fig 8) revealed presence of orthorhombic vaterite (ICSD File no. 98-011-5332) with peaks identified at 2θ values of 31.80°, 32.77° correlated with lattice indices (h k l) of (0 2 4) and (1 1 6) respectively.

**Fig 7:**
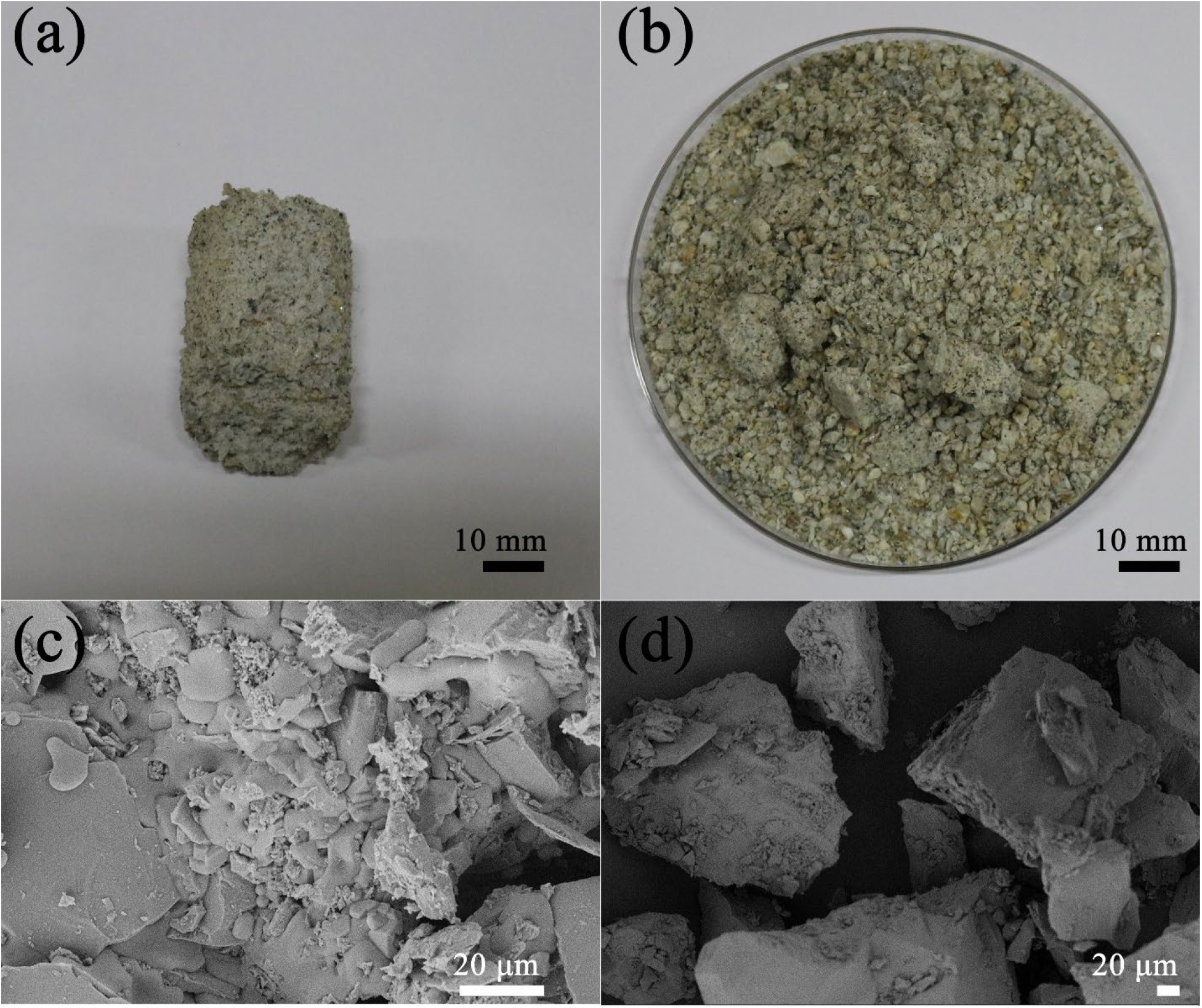
Bio-consolidation of sand (a) Photograph of bio-consolidated sample (b) photograph of the control sample with SMUC media (without bacteria) (c) SEM image of bio-consolidated sample, (d) SEM image of control sample.

**Fig 8:**
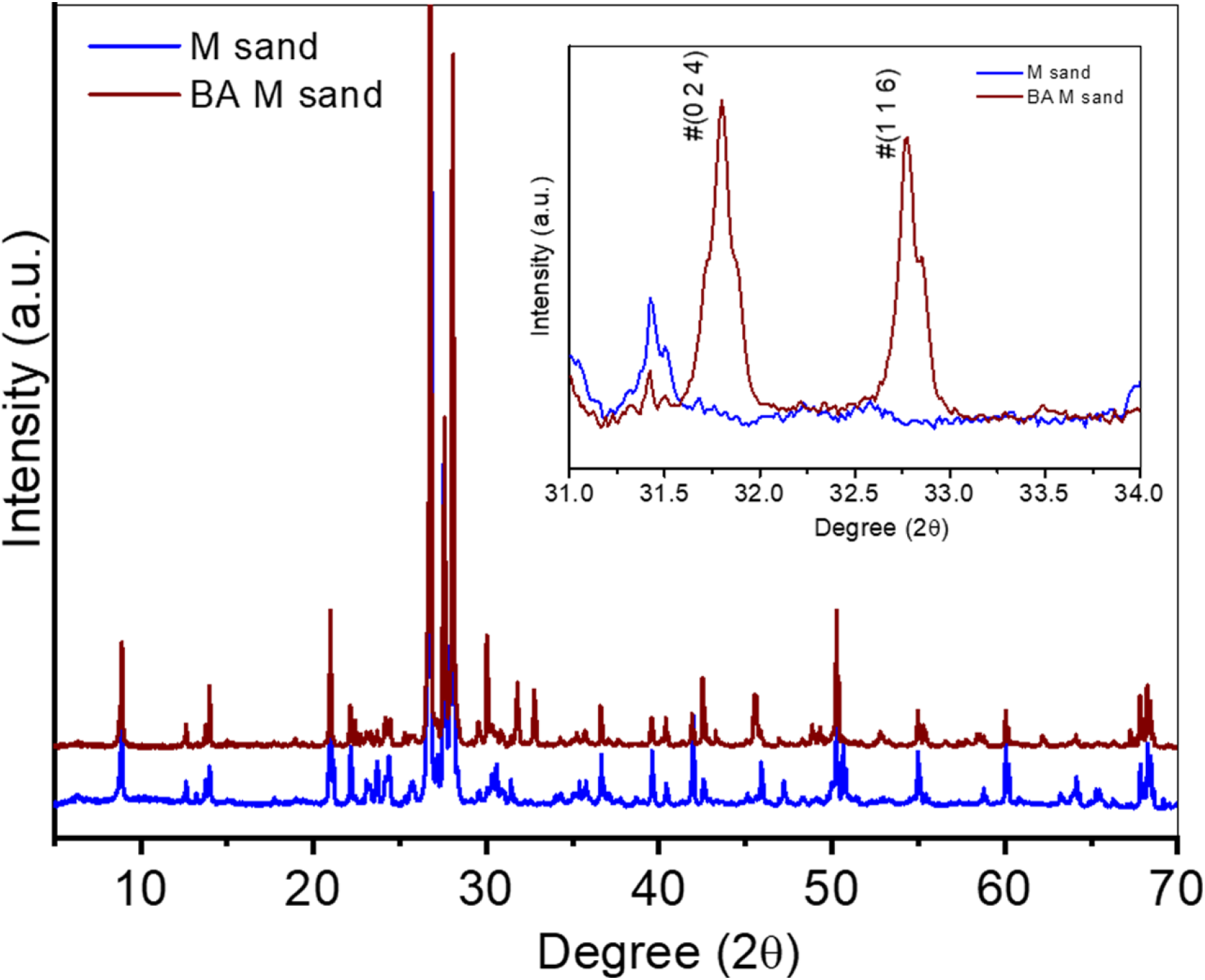
XRD pattern of bio-Consolidated sample, compared with sand control sample. # represents Vaterite – ICPD no. 98011532.

## 4. Discussion

The present study explored the novel urease producer soil bacterium for MICP application, which was identified as *Bacillus velezensis* through 16S rRNA gene sequencing. The important factor for MICP via ureolytic pathway is the selection of strain and its specificity based on the rate of urea hydrolysis. *B. velezensis* has shown potential for urea hydrolysis and degradation of urea was observed after 10 hrs of inoculation on urea agar plate which indicates its suitability for MICP mediated applications. Historically, *Bacillus velezensis* was isolated from the river of Velez in Ma ́laga (Southern Spain) and explored by Ruiz-Garcia et al.[32]. This novel isolated strain was characterized as a gram-positive bacterium that can tolerate a varied range of pH 5.0-10 and temperature 15 to 45°C.

Further, the ureolytic activity of isolated strain was explored for calcium carbonate precipitation under flask conditions with biopolymer (guar gum), cationic polypeptide (L-phenylalanine) additives and compared with media composition with no additives (SMUC). These naturally occurring polymer additives accelerated calcium carbonate precipitation, and a significant enhancement in the amount of precipitation was also observed. Different phases of calcium carbonate precipitation induced by bacterial activity have been reported in numerous studies [6, 16, 33-35]. Major forms of calcium carbonate precipitate are calcite, aragonite, and vaterite of which vaterite and calcite are the most commonly reported polymorphs of bacterial calcium carbonate precipitates [36]. Precipitation found in this study with newly isolated bacterium incubated medium was dominated by vaterite, which is a metastable form of calcium carbonate. SEM images also depicted significantly different precipitate morphology under different media supplementation. The role of these polymers as a media additive for MICP mediated consolidation of loose mass should be further investigated.

Guar gum gave promising results in terms of the amount of precipitation and calcite content which is known to be the stable most form of calcium carbonate [37]. The effect of guar gum additive on bacterial activity was extensively studied with four different concentrations by measuring change in the pH, ammonium concentration and total reducing sugar every 24 hours for 7 days. It was found that isolated strain was able to survive with different concentrations of guar gum, a complex biopolymer.

This biopolymer supplemented media has shown positive influence on calcite precipitation by enhancing the pH of the medium due to urea hydrolysis and release of ammonia. The experiment was initiated at an acidic pH of 5.5 because any adjustment in pH towards alkalinity would result in quick abiotic precipitation which is undesirable. Tanushree et al. [6] had also concluded that initial media pH played an important role in microbial induced calcite precipitation as when media pH was kept at 9, quick abiotic precipitation was observed. Microbial precipitation is influenced by pH of the medium because urease enzyme hydrolyses urea into ammonium ions, increases the pH and thus provides suitable micro-environment to proceed the reactions [6, 38, 39]. In the present study, maximum urea hydrolysis by bacterial activity was observed with higher concentrations of guar gum as evident from enhancement in medium pH and ammonium ion concentration.

Bio-degradation of complex polymer (i.e., guar gum) into small moiety was observed by *Bacillus velezensis* activity. In case of media without guar gum, as glucose is a monosaccharide which can metabolize easily, degrades till 24 hours and followed by its utilization by bacteria thereafter as is evident from total sugar content characteristics with time. With guar gum additive, net degradation by bacterial activity lasted up to 48 hours (in 0.25% guar gum) and 72 hours in rest of the cases followed by its utilization by bacterial strain. Polymer degradation releases free sugar which could be readily available in the growth medium for bacterial growth and resultant enhancement in the activity. It is reported that the organic matrix covering calcium carbonate precipitation is made up of either proteins or polysaccharides and these macromolecules play an important role in the synthesis of biominerals either by providing a structural framework, or regulating its entire dynamic process such as the nucleation site, growth of crystal and direction of orientation etc.[27, 40].

Bio consolidation mediated by *B. velezensis* exhibited the presence of vaterite. SEM images revealed sand particles and precipitates aggregated by covering of bacterial-induced matrix. Similar result was reported by Fadwa et al. [41] with MICP mediated consolidated gypsum plaster, which was covered with organics matrix along with meso-crystals of vaterite. Such organics matrix embedded with vaterite meso-crystals might be the reason behind the aggregation of the loose mass of the soil particles. Further experiments shall be performed to investigate the co-relation between organic matrix along with vaterite mediated consolidation.

## 5. Conclusion

In the present work, an efficient carbonate-precipitating soil bacterium was discovered and identified as *Bacillus velezensis*. The MICP characteristics of this strain were evaluated under three different media compositions. The isolated strain has given significant amount of calcite precipitation in all cases, with highest amount in case of guar gum additive as it was able to degrade guar as a biopolymer supplemented in the medium. It was further subjected to four different conditions of guar and maximum activity was observed in case of 1% guar. The strain was subjected to bio-cementation and has successfully consolidated locally available sand. This unexplored strain had shown potential towards MICP and bio-consolidation of loose mass. To the best of the authors’ knowledge, this is the first ever report on bio-consolidation mediated by *Bacillus velezensis* depicting positive influence of biopolymer on calcite precipitation. The encouraging results of this study indicate that, this locally available strain could be a model solution for the construction industry (e.g., bio-bricks by a green route) and other related MICP based applications.

## Acknowledgments

Authors would like to acknowledge Dr. Swetha Seshagiri, Associate Professor Center for Incubation Innovation Research and Consultancy, Jyothy Institute of Technology Bangalore-560082, Karnataka, India for kind gift of soil isolates. Authors also would like to acknowledge Ms. Harshita Patangia for her help in performing lab experiments. The authors acknowledge funding from Indian Space Research Organization (ISRO) and also funding from Indian Institute of Science.

